## Supplementary Table for "Direct lysis RT-qPCR of SARS-CoV-2 in cell culture supernatant allows for fast and accurate quantification of virus, opening a vast array of applications"

**Supplementary Information**

Table S1: Primer sequences

| Primer Pair | Forward primer | Reverse primer |
|---|---|---|
| **CDC N1** | GACCCCAAAATCAGCGAAAT | TCTGGTTACTGCCAGTTGAATCTG |
| **CDC N2** | TTACAAACATTGGCCGCAAA | GCGCGACATTCCGAAGAA |
| **CDC N3** | GGGAGCCTTGAATACACCAAAA | TGTAGCACGATTGCAGCATTG |
| **DZIF N** | CACATTGGCACCCGCAATC | GAGGAACGAGAAGAGGCTTG |
| **DZIF RdRp** | GTGARATGGTCATGTGTGGCGG | CARATGTTAAASACACTATTAGCATA |

Table S2: SARS-CoV-2 RNA template sequence for the DNA constructs (black) and the corresponding *in vitro* transcribed RNA (red/black)

| Template DNA / RNA | Gene | Sequence | Length (bases) | Primers |
|---|---|---|---|---|
| F10<br>tempRNA 10 | RdRp | gaatactcaagctatgcatcaagcttggtaccgagctcggatccactagtaacggccgccagtgtgctggaattcgcc<br>cgaatactcaagctatgcatcaagcttggtaccgagctcggatccactagtaacggccgccagtgtgctggaattcgc<br>cccaagtattgagtgaaatggtcatgtgtggcggttcactatatgttaaaccaggtggaacctcatcaggagatgccac<br>aactgcttatgctaatagtgtttttaacatttgtcaagctgtcacggccaatgttaatgcacttttatctactgatggtaacaa<br>aattgccgataagtatgtccgcaatttacaacacagactttatgagtgtctctatagaaatagagatgttgacacagactt<br>tgtgaatgagttttacgcatatttgcgtaaacatttctcaatgatgatactctctgacgatgctgttgtgtgtttcaatagcactt<br>atgcatctcaaggtctagtggctagcataaagaactttaagtcagttctttattatcaaaacaatgtttttatgtctgaagca<br>aaatgttggactgagactgaccttactaaaggacctcatgaattttgctctcaacatacaatgctagttaaacagggtga<br>tgattatgtgtaccttccttacccagatccatcaagaatcctaggggccggctgttttgtagatgatatcgtaaaaacaga<br>tggtacacttatgattgaacggttcgtgtctttagctatagatgcttacccacttactaaacatcctaatcaggagtatgctg<br>atgtctttcatttgtacttacaatacataagaaagctacatgatgagttaacaggacacatgttagacatgtattctgttatg<br>cttactaatgataacacttcaaggtattgggaacctgagttttatgaggctatgtacacaccgcatacagtcttacaggct<br>gttggggcttgtgttctttgcaattcacagacttcattaagatgtggtgcttgcatacgtagaccattcttatgttgtaaatgct<br>gttacgaccatgtcatatcaacatcacataaattagtcttgtctgttaatccgtatgtttgcaatgctccaggttgtgatgtca<br>cagatgtgactcaactttacttaggaggtatgagctattattgtaaatcacataaaccacccattagttttccattgtgtgct<br>aatggacaagtttttggtttatataaaaatacatgtgttggtagcgataatgttactgactttaatgcaattgcaacatgtga<br>ctggacaaatgctggtgattacattttagctaacacctgtactgaaagactcaagctttttgcagcagaaacgctcaaa<br>gctactgaggagacattgggcgaattctgcagatatccatcacactggcttaagggcgaattctgcagatatccatcac<br>actggc | 1361/<br>1476 | DZIF<br>RdRp |

| | | | | |
|---|---|---|---|---|
| F18<br>tempRNA 18 | N | gaatactcaagctatgcatcaagcttggtaccgagctcggatccactagtaacggccgccagtgtgctggaattcgcc<br>cgaatactcaagctatgcatcaagcttggtaccgagctcggatccactagtaacggccgccagtgtgctggaattcgc<br>cctgccagatcagtttcacctaaactgttcatcagacaagaggaagttcaagaactttactctccaatttttcttattgttgc<br>ggcaatagtgtttataacactttgcttcacactcaaaagaaagacagaatgattgaactttcattaattgacttctatttgtg<br>ctttttagcctttctgctattccttgttttaattatgcttattatcttttggttctcacttgaactgcaagatcataatgaaacttgtca<br>cgcctaaacgaacatgaaatttcttgttttcttaggaatcatcacaactgtagctgcatttcaccaagaatgtagtttacag<br>tcatgtactcaacatcaaccatatgtagttgatgacccgtgtcctattcacttctattctaaatggtatattagagtaggagc<br>tagaaaatcagcacctttaattgaattgtgcgtggatgaggctggttctaaatcacccattcagtacatcgatatcggtaa<br>ttatacagtttcctgtttaccttttacaattaattgccaggaacctaaattgggtagtcttgtagtgcgttgttcgttctatgaag<br>actttttagagtatcatgacgttcgtgttgttttagatttcatctaaacgaacaaactaaaatgtctgataatggaccccaaa<br>atcagcgaaatgcaccccgcattacgtttggtggaccctcagattcaactggcagtaaccagaatggagaacgcagt<br>ggggcgcgatcaaaacaacgtcggccccaaggtttacccaataatactgcgtcttggttcaccgctctcactcaacat<br>ggcaaggaagaccttaaattccctcgaggacaaggcgttccaattaacaccaatagcagtccagatgaccaaattg<br>gctactaccgaagagctaccagacgaattcgtggtggtgacggtaaaatgaaagatctcagtccaagatggtatttct<br>actacctaggaactgggccagaagctggacttccctatggtgctaacaaagacggcatcatatgggttgcaactgag<br>ggagccttgaatacaccaaaagatcacattggcacccgcaatcctgctaacaatgctgcaatcgtgctacaacttcct<br>caaggaacaacattgccaaaaggcttctacgcagaagggagcagaggcggcagtcaagcctcttctcgttcctcat<br>cacgtagtcgcaacagttcaagaaattcaactccaggcagcagtaggggaacttctcctgctagaatggctggcaat<br>ggcggtgatgctgctcttgctttgctgctgcttgacagattgaaccagcttgagagcaaaatgtctggtaaaggccaac<br>aacaacaaggccaaactgtcactaagaaatctgctgctgaggcttctaagaagcctcggcaaaaacgtactgccac<br>taaagcatacaatgtaacacaagctttcggcagacgtggtccagaacaaacccaaggaagggcgaattctgcaga<br>tatccatcacactggcttaagggcgaattctgcagatatccatcacactggc | 1611/<br>1726 | CDC N1<br>CDC N3<br>DZIF N |
| F19<br>tempRNA 19 | N | gaatactcaagctatgcatcaagcttggtaccgagctcggatccactagtaacggccgccagtgtgctggaattcgcc<br>cgcgaattgggccctctagatgcatgctcgagcggccgccagtgtgatggatatctgcagaattcgccccagaacaa<br>acccaaggaaattttggggaccaggaactaatcagacaaggaactgattacaaacattggccgcaaattgcacaat<br>ttgcccccagcgcttcagcgttcttcggaatgtcgcgcattggcatggaagtcacaccttcgggaacgtggttgacctac<br>acaggtgccatcaaattggatgacaaagatccaaatttcaaagatcaagtcattttgctgaataagcatattgacgcat<br>acaaaacattcccaccaacagagcctaaaaaggacaaaaagaagaaggctgatgaaactcaagccttaccgca<br>gagacagaagaaacagcaaactgtgactcttcttcctgctgcagatttggatgatttctccaaacaattgcaacaatcc<br>atgagcagtgctgactcaactcaggcctaaactcatgcagaccacacaaggcagatgggctatataaacgttttcgct<br>tttccgtttacgatatatagtctactcgggcgaattccagcacactggcggccgttactagtggatccgttaagggcgaat<br>tctgcagatatccatcacactggc | 610/<br>725 | CDC N2 |

6
